# KDM5-mediated activation of genes required for mitochondrial biology is necessary for viability in *Drosophila*

**DOI:** 10.1101/2023.05.23.541787

**Authors:** Michael F Rogers, Owen J Marshall, Julie Secombe

## Abstract

The precise coordination of gene expression is critical for developmental programs, and histone modifying proteins play important, conserved roles in fine-tuning transcription for these processes. One such family of proteins are KDM5 enzymes that interact with chromatin through demethylating H3K4me3 as well as demethylase-independent mechanisms that remain less understood. The single *kdm5* ortholog in *Drosophila* is an essential gene that has crucial developmental roles in a neuroendocrine tissue, the prothoracic gland. To characterize the regulatory functions of KDM5, we examined its role in coordinating gene expression programs critical to cellular homeostasis and organismal viability in larval prothoracic gland cells. Utilizing targeted genetic experiments, we analyzed the relationship between critical cell signaling pathways, particularly MAPK, and the lethality caused by loss of *kdm5*. Integrating KDM5 genome binding and transcriptomic data revealed conserved and tissue-specific transcriptional programs regulated by KDM5. These experiments highlighted a role for KDM5 in regulating the expression of a set of genes critical for the function and maintenance of mitochondria. This gene expression program is key to the essential functions of KDM5, as expression of the mitochondrial biogenesis transcription factor Ets97D/Delg, the *Drosophila* homolog of GABPα, in prothoracic gland cells suppressed the lethality of *kdm5* null animals. Consistent with this, we observed morphological changes to mitochondria in the prothoracic gland of *kdm5* null mutant animals. Together, these data establish KDM5-mediated cellular functions that are both important for normal development and could also contribute to KDM5-linked disorders when dysregulated.

## INTRODUCTION

Transcriptional regulators function as powerful gatekeepers that enable cells to access and utilize the information stored in the genome. The dynamics of chromatin organization and transcriptional mechanisms must therefore be carefully coordinated to orchestrate the gene expression programs required for proper development. Conversely, improper function of transcriptional regulators can underlie the defective cellular processes that lead to dysfunction and disease (Mirabella et al., 2016, Lee and Young, 2013). Within this realm of biology, chromatin-modifying proteins interface with histone protein tails through writing, reading, and erasing post-translational modifications to organize gene expression. KDM5 (<u>Lysine D</u>e<u>m</u>ethylase <u>5</u>) proteins are one such family of chromatin-modifiers that are named for their ability to remove trimethylation of lysine 4 on histone H3 (H3K4me3), a mark generally found near the transcriptional start sites of actively expressed genes (Chan et al., 2022).

Mammalian cells encode four paralogous KDM5 proteins: KDM5A, KDM5B, KDM5C, and KDM5D. The importance of gene regulation by KDM5 family proteins is demonstrated by their links to human disorders. All four KDM5 genes have been observed to show altered expression across a variety of cancer types, of which breast and prostate cancer are the most well characterized (Ohguchi and Ohguchi, 2022, Blair et al., 2011). The relationship between KDM5A, KDM5B, and tumorigenesis appears to be primarily oncogenic, with a range of cancers showing increased expression of either of these two paralogs. Rather than being linked to the regulation of a single process in malignant cells, KDM5A and KDM5B contribute to many facets of tumorigenesis including the regulation of genes linked to cell cycle control, DNA repair and angiogenesis (Yoo et al., 2022, Ohguchi et al., 2021, Taylor-Papadimitriou and Burchell, 2022, Ohguchi and Ohguchi, 2022). The roles of KDM5C and KDM5D in malignancies are less defined, although, in contrast to KDM5A and KDM5B, it is generally reduction of these proteins that is observed in various cancers, most notably renal carcinomas (Tricarico et al., 2020). The genetic association between KDM5 proteins and neurodevelopmental disorders, including intellectual disability and autism spectrum disorders, is more clearly caused by loss of function variants in *KDM5A*, *KDM5B* or *KDM5C* (Hatch and Secombe, 2021, Yoo et al., 2022). Consistent with this, mouse and cell culture models have shown that Kdm5A, Kdm5B and/or Kdm5C are needed for proper neuronal differentiation and morphology (Iwase et al., 2017, Iwase et al., 2016, Harrington et al., 2022, El Hayek et al., 2020). However, while KDM5 proteins are clearly required for normal brain function, the transcriptional programs critical for typical cognitive development remain unknown. It also remains unclear whether similar or distinct transcriptional programs etiologically link KDM5 to malignancies and to brain development. In this regard, it is notable that although cancer and intellectual disability have vastly different clinical manifestations, alterations in the activity of other regulatory factors, such as members of the MAPK/Ras (mitogen-activated protein kinase) and PI3K signaling cascades, are also linked to these same two disorders (Borrie et al., 2017). Thus, it remains possible that dysregulation of overlapping pathways contributes to both altered cognition and tumorigenesis.

Defining precisely how changes to KDM5 protein function leads to cancer or intellectual disability would be greatly facilitated by efforts to understand the fundamental transcriptional activities of KDM5 proteins. To date, most attempts to define these links have focused on the canonical histone demethylase activity. However, it is becoming increasingly apparent that KDM5 and other chromatin-modifying proteins also perform important non-catalytic gene regulatory functions (Ohguchi and Ohguchi, 2022, Paroni et al., 2018, Cao et al., 2014, Aubert et al., 2019, Morgan and Shilatifard, 2023). Demethylase-dependent and independent activities of KDM5 proteins have been shown to play roles in both cancer and intellectual disability (Iwase et al., 2007, Vallianatos et al., 2018, Paroni et al., 2018). This is also true in the animal model *Drosophila melanogaster*, which encodes a single KDM5 protein that is likely to function incorporating activities of all four mammalian paralogs. Establishing its critical role in developmental processes is the fact that a null allele in *Drosophila kdm5* (*kdm5^140^*) causes lethality during development (Drelon et al., 2018). The essential functions of KDM5 are independent of its enzymatic demethylase function, however, as animals harboring loss-of-function mutations in the enzymatic Jumonji C (JmjC) domain survive to adulthood (Drelon et al., 2018, Li et al., 2010). Characterizing the role of KDM5 during *Drosophila* development therefore provides an opportunity to uncover new pathways and gene-regulatory mechanisms that will expand our understanding of this family of multi-domain proteins.

Several cell types in *Drosophila* have been shown to require KDM5 during development. Consistent with the established link between genetic variants in human *KDM5* genes and intellectual disability, KDM5 is necessary for proper neuronal development and functioning (Belalcazar et al., 2021, Hatch et al., 2021, Zamurrad et al., 2018). However, these neuronal activities of KDM5 are not necessarily involved in its essential developmental functions, as restoring *kdm5* expression pan-neuronally does not rescue lethality (Drelon et al., 2019). KDM5 has also been linked to immune function in larval hemocytes, but, in a similar manner to neurons, this cell type does not account for its essential activities (Drelon et al., 2019, Moran et al., 2015). The only single tissue in which re-expression of *kdm5* is sufficient to rescue lethality is the prothoracic gland (Drelon et al., 2019). *kdm5^140^* (null) animals rescued by prothoracic gland-specific *kdm5* expression develop into adult flies, however, they survive at a lower frequency than animals expressing *kdm5* ubiquitously, which indicates that KDM5 also has essential developmental functions in other tissues. Nevertheless, this rescue demonstrates that within prothoracic gland cells, KDM5 regulates the expression of genes crucial to proper organismal development.

A neuroendocrine tissue, the prothoracic gland serves as a master coordinator of numerous intracellular cellular processes, tissue growth, and organismal transitions that are essential to development through its production of the steroid hormone ecdysone (Kamiyama and Niwa, 2022, Texada et al., 2020). This tissue is also a well-established model for understanding how key signaling pathways are integrated to govern hormone dynamics and animal maturation, including the MAPK, Salvador-Warts-Hippo-Yorkie (SHW), target of rapamycin (TOR) and insulin and insulin-like signaling (IIS) cascades. These pathways are known to converge on cellular processes, such as autophagy, that are critical for regulating metabolism and hormone production in the prothoracic gland (Texada et al., 2019, Nagata et al., 2022, Moeller et al., 2017). Like KDM5 family proteins, the dysregulation of these pathways is implicated in human disorders including cancer and neurodevelopmental disorders (Vithayathil et al., 2018, Kim and Choi, 2010, Zanconato et al., 2016, Tian et al., 2019, Williamson et al., 2014). Studying the functions of KDM5 in the prothoracic gland will therefore provide important information about the relationship between KDM5-regulated gene expression and these critical pathways.

KDM5 plays at least two distinct roles in cells of the prothoracic gland. The first is in the regulation of larval growth rate. Although *kdm5* null mutants can eventually reach wild-type size and undergo metamorphosis, they take as much as twice the amount of time to progress through the stages of development and exhibit corresponding reduced ecdysone levels (Drelon et al., 2018). In this context, KDM5 promotes the endoreplication of prothoracic gland cells, which increases ploidy in order to maximally express ecdysone biosynthetic factors (Drelon et al., 2019, Ohhara et al., 2017, Ohhara et al., 2019). The role of KDM5 in facilitating normal growth rate, however, is separate to its role in survival, as restoring normal developmental timing to *kdm5* mutant animals does not alter their viability. The role of KDM5 in promoting animal survival does involve the MAPK signaling pathway, as *kdm5* null mutant animals show decreased MAPK signaling and activating this pathway in the prothoracic gland suppresses *kdm5* mutant lethality. However, whether this effect is specific to the MAPK pathway, and which downstream cellular processes link KDM5, MAPK, and viability, remain to be established.

Here we examine KDM5 function in the prothoracic gland as a means to broadly understand how this chromatin modifier regulates critical cellular processes. Extending from our previous studies, we explore the role of the MAPK and parallel pathways in mediating the lethality caused by loss of *kdm5* by taking targeted approaches based on the known signaling cascades. We additionally take unbiased approaches to define the transcriptional targets of KDM5. Among these targets, we identified mitochondrial biology as a candidate process for which KDM5-mediated regulation could play critical roles during development. Reinforcing these connections, the lethality of the *kdm5* null allele can be suppressed by expression of Ets97D/Delg, the *Drosophila* homolog of GA Binding Protein Transcription Factor Subunit Alpha (GABPα), a known activator of genes necessary for cellular respiration. Supporting this, prothoracic gland cells of *kdm5* mutant animals show altered mitochondrial morphology dynamics. Together, this study provides new insights into the link between KDM5-regulated transcription, mitochondrial function, and vital cellular processes needed to coordinate development.

## RESULTS

### Activating MAPK signaling suppresses *kdm5* null lethality independently of autophagy regulation

To better understand the critical developmental roles of KDM5, we first sought to further investigate the link between *kdm5*-induced lethality and activation of MAPK signaling (Drelon et al., 2019). From yeast to humans, the MAPK signaling cascade is used by cells to regulate a myriad of downstream cellular events in a context-dependent manner (Widmann et al., 1999, Yang et al., 2013, Eblen, 2018, Pan and O’Connor, 2021). In the prothoracic gland of *Drosophila*, the MAPK pathway is one of several signaling networks that regulates ecdysone biosynthesis (Fig. 1A). To further characterize the relationship between KDM5 and MAPK, we took a candidate-based approach, testing upstream and downstream components of this cascade for an effect on *kdm5*-induced lethality (Fig. 1B). We used spookier-Gal4 (*spok*-Gal4) to drive expression of transgenes in a tissue-specific manner within the prothoracic gland, hereafter written as “*spok*>*transgene*” (Fig. 1C) (Drelon et al., 2019, Shimell et al., 2018, Pan and O’Connor, 2021). As quantified previously, the ability of tested transgenes to mediate *kdm5^140^* (null allele) survival into adulthood was calculated, and for these experiments, this survival index was normalized to that observed by *spok*-Gal4-driven expression of KDM5 (% *spok*>*kdm5*, see Methods).

**Figure 1:**
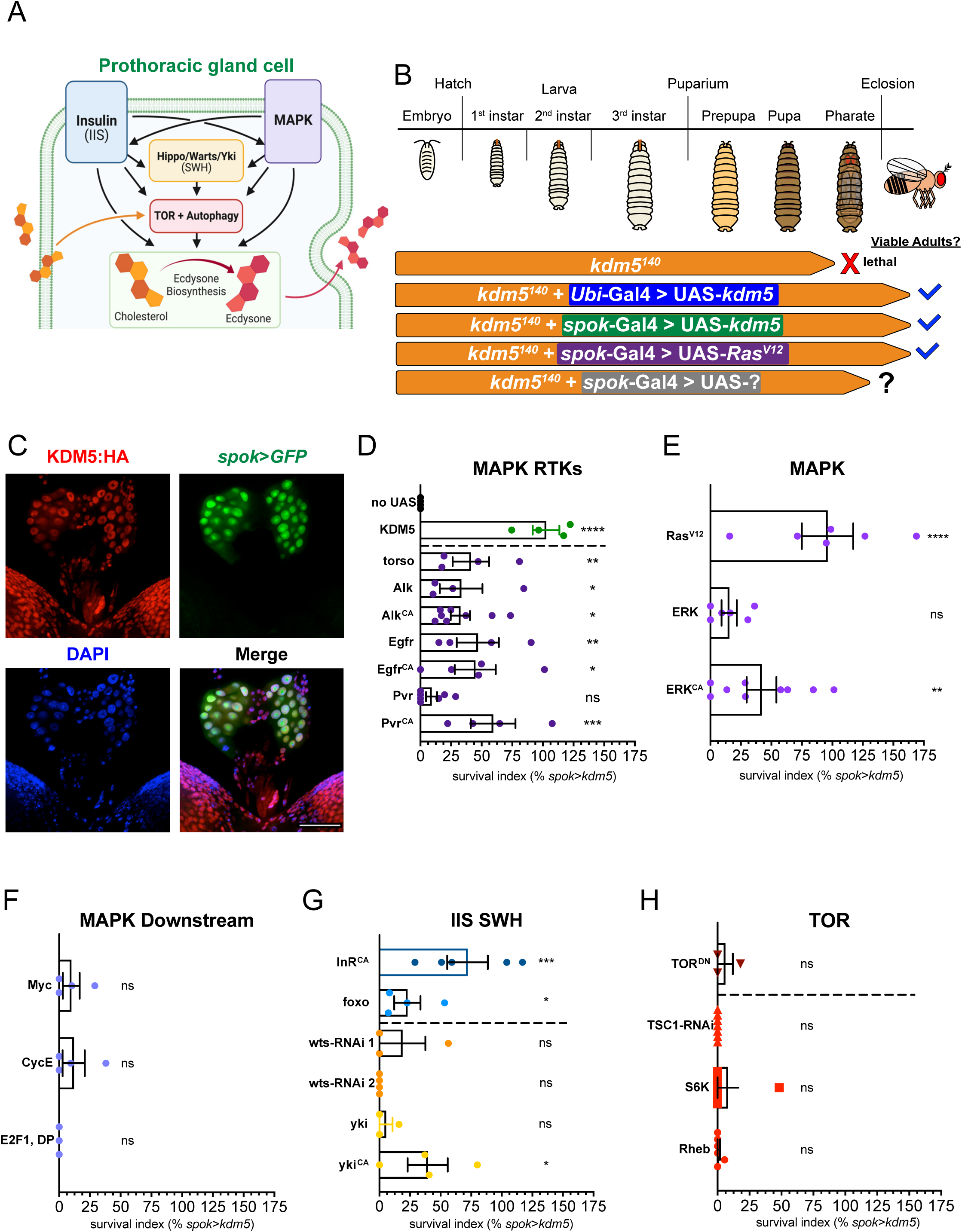
MAPK signaling robustly suppresses *kdm5^140^* lethality independent of autophagy regulation. (A) Schematic summarizing major cellular signaling pathways known to regulate prothoracic gland cell function. Potential crosstalk interactions and common targets between pathways indicated by black arrows. Created with BioRender. (B) Schematic summarizing previous findings from Drelon et al. (2019) of *kdm5^140^* pupal pharate lethality suppression by transgene expression, including MAPK signaling via RasV^12^. (C) Maximum intensity Z-projection image of brain-ring gland complex of wandering L3 larva shows expression of endogenously-tagged KDM5:HA in the nuclei of the prothoracic gland. Prothoracic gland marked by *spok*-Gal4-driven GFP expression and nuclei marked by DAPI stain. Scale bars represent 50 μm. (D) Quantification of survival index for expression of MAPK-activating RTKs in *kdm5^140^*background relative to *spok*-Gal4>UAS-*kdm5* (green data points). n = 191-722 (mean n = 509) per genotype tested. ****p<0.0001, ***p<0.001, **p<0.01, *p<0.05; ns, not significant (Fisher’s exact test compared to no UAS control (black data points)). Error bars: mean + s.e.m. (E) Quantification of survival index for expression of MAPK signaling components in *kdm5^140^* background relative to *spok*-Gal4>UAS-*kdm5*. n = 467-800 (mean n = 627) per genotype tested. ****p<0.0001, **p<0.01; ns, not significant (Fisher’s exact test compared to no UAS control). Error bars: mean + s.e.m. (F) Quantification of survival index for expression of candidate factors downstream of MAPK in *kdm5^140^* background relative to *spok*-Gal4>UAS-*kdm5*. n = 283-484 (mean n = 378) per genotype tested. ns, not significant (Fisher’s exact test compared to no UAS control). Error bars: mean + s.e.m. (G) Quantification of survival index for expression of IIS (insulin and insulin-like signaling) and SWH (Salvador-Warts-Hippo-Yorkie) signaling components in *kdm5^140^* background relative to *spok*-Gal4>UAS-*kdm5*. n = 292-921 (mean n = 466) per genotype tested. ***p<0.001, *p<0.05; ns, not significant (Fisher’s exact test compared to no UAS control). Error bars: mean + s.e.m. (H) Quantification of survival index for expression of autophagy-regulating components of TOR signaling in *kdm5^140^* background relative to *spok*-Gal4>UAS-*kdm5*. n = 352-926 (mean n = 638) per genotype tested. ns, not significant (Fisher’s exact test compared to no UAS control). Error bars: mean + s.e.m.

Based on the suppression of *kdm5^140^* lethality by expression of the receptor tyrosine kinase (RTK) Torso and activated Ras (Ras^V12^) in Drelon et al. (2019), we tested whether other RTKs upstream of MAPK, or whether the downstream kinase ERK, could restore *kdm5^140^* viability (Drelon et al., 2019). In parallel to Torso, which receives neuronal stimulation via the prothoracicotropic hormone (PTTH) neurotransmitter, the anaplastic lymphoma kinase (Alk), epidermal growth factor receptor (Egfr), and PDGR and VEGF-receptor related (Pvr) RTKs can also activate MAPK signaling and impact ecdysone biosynthesis in response to extracellular inputs (Pan and O’Connor, 2021, Cruz et al., 2020). *spok*-Gal4-driven expression of wild-type or constitutively active (^CA^) forms of each of these receptors resulted in partial suppression of lethality with a mean survival index of 33.2% (Fig. 1D). Likewise, *spok*>*erk* and *spok*>*erk^CA^* resulted in survival indices of 17.4% and 41.8%, respectively (Fig. 1E). While most MAPK transgenes tested significantly restored *kdm5^140^* viability, none were as effective as Ras^V12^, which had an average survival index of 78.1% (Fig. 1E). This possibly reflects stronger activation of signaling by the Ras^V12^ transgene, particularly due to the role of post-translational modifications in regulating the MAPK cascade. Similar to the rescue of *kdm5^140^* by expression of KDM5 in the prothoracic gland, adult flies obtained through expression of RTKs or ERK had successfully formed adult structures but with an outstretched wings phenotype and reduced lifespan (Supp Fig. 1) (Drelon et al., 2019). Combined, these data confirm that augmenting MAPK signaling through various means of activation, not only through the Torso-Ras axis, all restore *kdm5^140^* viability. The downstream effectors that mediate MAPK signaling in the prothoracic gland remain unknown; however, in other contexts, regulatory proteins such as Myc, the E2F1/DP heterodimer, and cell cycle mediators can be regulated by this cascade (Zhang and Liu, 2002). Because these specific transcription factors and cellular processes have also been associated with mammalian or *Drosophila* KDM5 function in other contexts, we next tested their ability to suppress *kdm5^140^* lethality (Secombe et al., 2007, Benevolenskaya et al., 2005, Drelon et al., 2019). Expression of Myc, E2F1 and DP, or Cyclin E did not alter *kdm5*-induced lethality, however, suggesting that other, unidentified regulators of gene expression function with KDM5 in the context of the prothoracic gland (Fig. 1F).

To determine whether this relationship with *kdm5^140^*lethality is specific to the MAPK pathway, we examined other signaling pathways that mediate prothoracic gland function, many of which show extensive crosstalk (Fig. 1A). Specifically, we tested the insulin and insulin-like growth factor signaling (IIS), Salvador-Warts-Hippo-Yorkie (SWH), and the target of rapamycin (TOR) pathways. These three pathways are among the best characterized to date for their roles in the prothoracic gland, particularly in the regulation of homeostatic metabolic processes such as autophagy and lipid processing for hormone production (Texada et al., 2019, Danielsen et al., 2013, Danielsen et al., 2014, Danielsen et al., 2016). To test the IIS cascade, we expressed an activated form of the insulin receptor (*spok*>*InR^CA^*) or the downstream transcription factor foxo (*spok*>*foxo*). Expression of InR or foxo did result in suppression of *kdm5^140^* lethality with survival indices of 65.6% and 24.8%, respectively (Fig. 1G). Though we previously saw no defective activation of the IIS pathway by phoso-Akt via Western blot in *kdm5^140^* animals in Drelon et al. (2019), it is possible that ectopic insulin signaling activation can act on similar downstream targets or compensate in some other way for MAPK defects (Drelon et al., 2019). In contrast, SWH signaling, activated by expression of RNAi against Wts (*spok*>*wts-RNAi* #1 and 2) or overexpression of wild-type or constitutively active yki transgenes (*spok*>*yki*, *spok*>*yki^CA^*) did not consistently suppress lethality (Fig. 1G). For this pathway, suppression was limited to yki^CA^, which, similar to the IIS cascade, may indicate that activation of these signaling pathways is able to compensate for *kdm5^140^* MAPK defects. These results could be due to crosstalk between these pathways and/or upregulation of common inputs involved in regulation of ecdysone biosynthesis and prothoracic gland function.

Additionally, prothoracic gland cells have distinct energetic and other cellular homeostatic requirements due to their status as terminally differentiated and large polyploid cells, and therefore proper balance of TOR signaling has been shown to be critical for tissue function (Danielsen et al., 2016, Texada et al., 2019, Pan et al., 2019, Yamanaka, 2021, Pan et al., 2020). For this reason, we tested several manipulations of TOR signaling and autophagy via both activation (*spok*>*Rheb*, *spok*>*S6K*, *spok*>*TSC-RNAi*) and repression (*spok*>*TOR^DN^*). Interestingly, none of these TOR pathway manipulations affected *kdm5^140^* lethality (Fig. 1H). Thus, while regulation of autophagy is one cellular process on which all of these signaling pathways are known to converge, the lethality of *kdm5^140^* mutants does not appear to be from lack of TOR pathway regulation. Taken together, there appear to be multiple pathways capable of suppressing *kdm5^140^* lethality via activity in the prothoracic gland, but it is not yet clear whether these results are due to crosstalk between pathways or compensatory activation of shared downstream targets. Moreover, it remains an open question which downstream transcription factors are responsible for the cellular programs activated by this signaling that are crucial for development and adult viability.

### *kdm5* expression is required during mid to late larval stages for viability

Our candidate approaches identified regulatory pathways, but not key KDM5-mediated downstream processes linked to viability. We therefore performed transcriptomic and genomic-binding studies to investigate KDM5 function in an unbiased manner. Prior to carrying out these molecular studies, we first needed to determine the periods during development in which KDM5 is required. To do this, we ubiquitously expressed the UAS-*kdm5* transgene using *Ubi*-Gal4 within defined windows of time during development in the *kdm5^140^* background (Fig. 2A). To facilitate temporal activation of *kdm5* expression, we included a transgene ubiquitously expressing temperature-sensitive Gal80^ts^ (tub-Gal80^ts^) (McGuire et al., 2003). At 18°C, the Gal80^ts^ prevents UAS-*kdm5* transgene activation, thus *kdm5^140^* animals with tub-Gal80^ts^, *Ubi*-Gal4, and UAS-*kdm5* incubated at 18°C fail to reach adulthood (Fig. 2A). At 29°C, Gal80^ts^ is inactivated, which allows constitutive expression of the UAS-*kdm5* transgene and results in adult fly viability (Fig. 2A). At the permissive temperature of 29°C, we observe protein levels similar to both endogenous KDM5 and to our previously published system in which flies were grown at 25°C without Gal80^ts^ (Fig. 2B) (Drelon et al., 2019). Using this system, *kdm5* expression was turned on at progressively later days during development by transferring the animals from 18°C to 29°C (modeled in Fig. 2A). The extent to which temporally-restricted expression of *kdm5* rescued viability is reported as a survival index normalized to the rescue observed by continuous expression of *kdm5* (*Ubi*>*kdm5* at constant 29°C, see Methods). Temperature shifting animals early in development led to robust rescue (Fig. 2C). In contrast, activating the UAS-*kdm5* transgene in animals that had reached mid larval stages (2^nd^-3^rd^ instar) or later resulted in a failure to rescue adult viability (Fig. 2C). Thus, *kdm5* expression is required prior to pupal stages and as early as mid to late larval stages, although we cannot yet rule out additional roles later in development. Additional complementary experiments in which UAS-*kdm5* transgene expression was inhibited progressively later in development were also performed by shifting animals from 29°C to 18°C (modeled in Fig. 2A). These data revealed that transferring animals that had reached mid larval stages (2^nd^-3^rd^ instar) or earlier failed to robustly rescue viability, confirming key role(s) for KDM5 during the mid to late larval window of the *Drosophila* life cycle (Fig. 2D). Based on this temporal rescue data, we focused subsequent experiments of KDM5 function during the late larval development.

**Figure 2:**
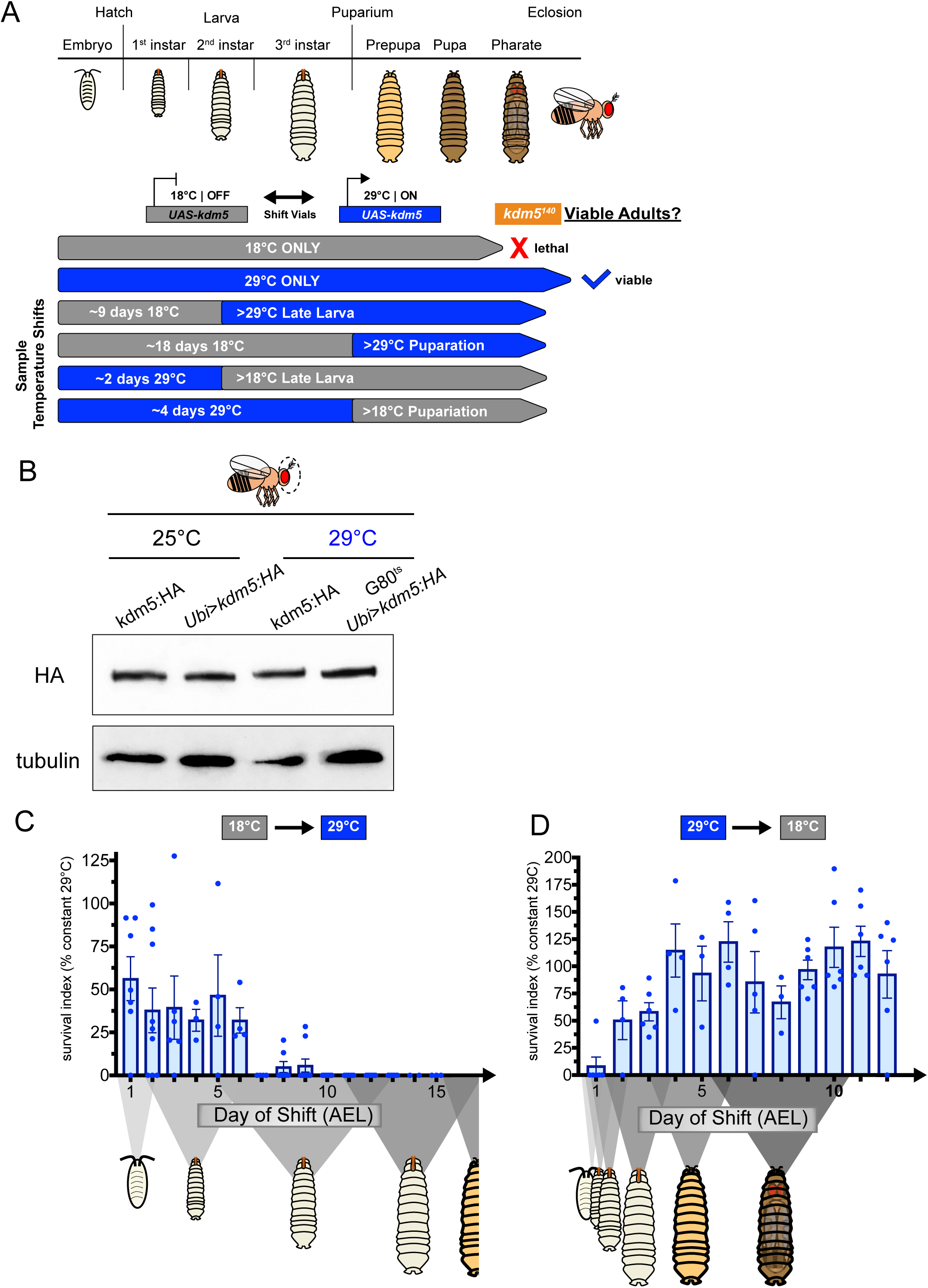
Temporally-restricted rescue KDM5 expression reveals requirements for KDM5 in mid-to-late larval stages. (A) Schematic demonstrating vial shifts between restrictive (18°C) and permissive (29°C) temperatures to constrain rescue KDM5 expression within defined developmental windows. (B) Western blot of adult heads showing comparable KDM5:HA protein levels (top) across control (kdm5:3xHA or *Ubi*>*kdm5:HA* (*kdm5^140^* background)) and temporal experiment (G80^ts^ *Ubi*>*kdm5:H*A (*kdm5^140^* background)) animals at standard (25°C) and experimental (29°C) temperatures. α-tubulin loading control. (C) Quantification of survival index for induction of expression of KDM5 at progressively later days during development (18°C to 29°C) in tub-Gal80^ts^ / +; *kdm5^140^*, *Ubi*-Gal4 / *kdm5^140^*; UAS*-kdm5:HA* / + animals relative to that of control vials kept at constant 29°C. X-axis schematic demonstrates developmental progression of *kdm5^140^* animals at 18°C at each day after egg lay (AEL). n = 101-275 (mean n = 153) per genotype tested. Error bars: mean + s.e.m. (D) Quantification of survival index for inhibition of expression of KDM5 at progressively earlier days during development (29°C to 18°C) in tub-Gal80^ts^ / +; *kdm5^140^*, *Ubi*-Gal4 / *kdm5^140^*; UAS*-kdm5:HA* / + animals relative to that of control vials kept at constant 29°C. X-axis schematic demonstrates developmental progression of *kdm5^140^* animals at 29°C at each day after egg lay (AEL). n = 103-210 (mean n = 128) per genotype tested. Error bars: mean + s.e.m.

### KDM5 directly regulates transcription of metabolic processes in the prothoracic gland

To investigate the roles of KDM5 in regulating gene expression programs within the prothoracic gland, we identified genomic regions bound by KDM5 in this tissue. Traditional genomic binding approaches such as ChIP-seq are limited for this small tissue that is comprised of ~50 cells. We therefore performed Targeted DamID (TaDa), which requires less input material and can be carried out with tissue and temporal-specific resolution, to survey the genomic targets of KDM5 in these cells (Hatch et al., 2021, Marshall and Brand, 2015, Marshall and Brand, 2017, Marshall et al., 2016a). We used *spok*-Gal4 to drive expression of a UAS transgene encoding a Dam:KDM5 fusion protein (or UAS-*dam* as the normalization control) exclusively in the prothoracic gland cells of wild-type animals. Using tub-Gal80^ts^, expression of Dam:KDM5 was restricted to the KDM5-critical late larval stages by shifting animals from 18°C to 29°C and collecting wander 3^rd^ instar larvae (120-168 hours after egg laying (AEL) at 18°C) (Fig. 3A). Confirming the robustness of our data, quadruplicate TaDa replicates showed a very strong correlation, and an average Dam:KDM5 binding profile was used for subsequent analyses (Supp Fig. 2). Similar to prior studies of KDM5 family proteins across species, a majority of KDM5 binding occurred within the proximity of promoter regions, particularly at nucleosomes bordering transcriptional start sites (TSS) (Fig. 3B, C) (Hatch et al., 2021, Liu and Secombe, 2015, Lloret-Llinares et al., 2012, Beshiri et al., 2012, Iwase et al., 2016, Wang et al., 2023). This localization at or near promoters enabled us to unambiguously identify nearby genes as candidate targets of KDM5 regulation.

**Figure 3:**
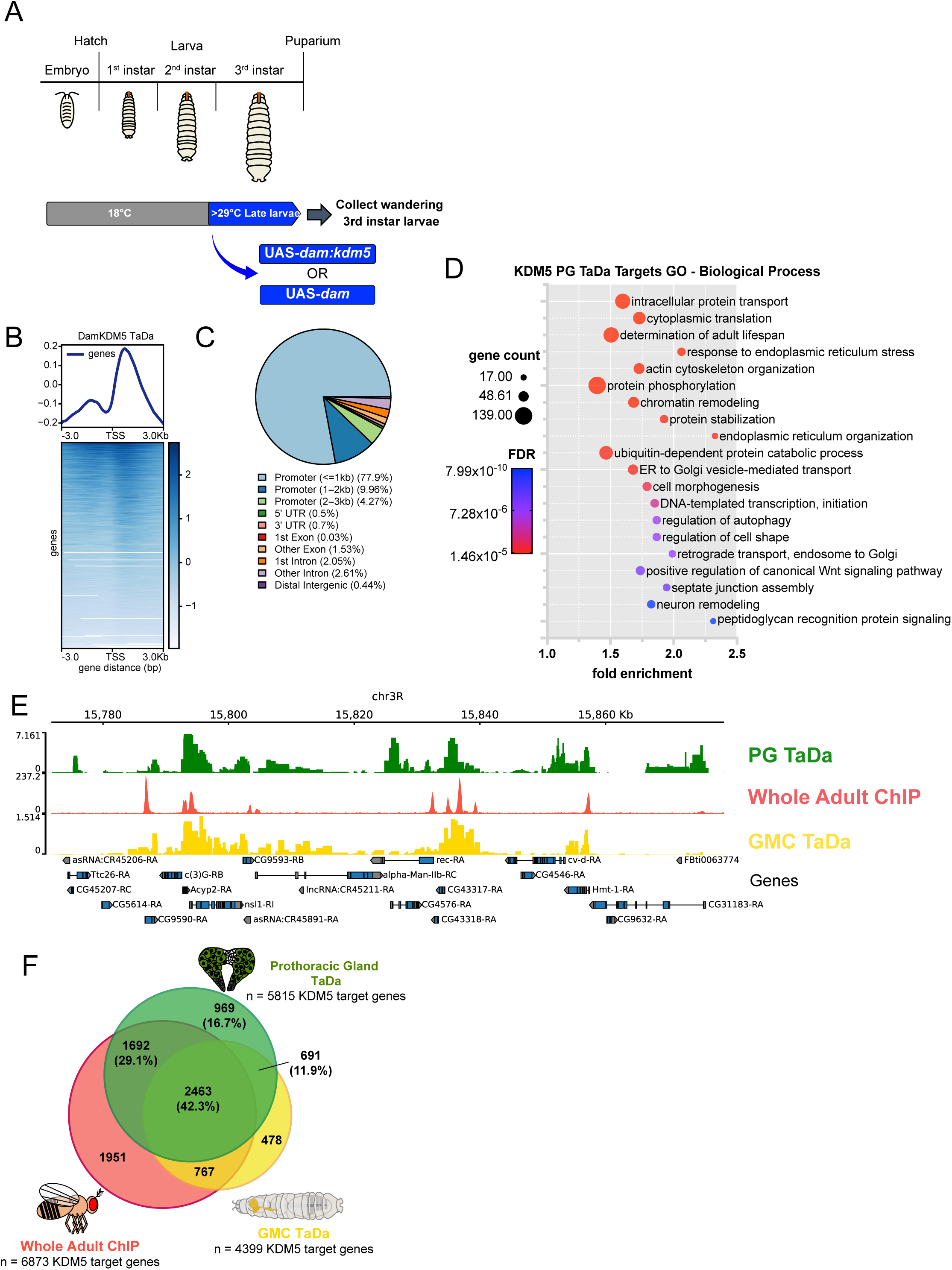
Genome binding profiling of KDM5 by targeted DamID identifies both conserved and tissue-specific target genes. (A) Schematic demonstrating time course of Targeted DamID (TaDa) experiment, which restricted *spok*-Gal4-driven transgene expression (UAS-*dam:kdm5* or UAS-*dam*) to last 48 hours of larval development. The TaDa experiment was performed in quadruplicate for each genotype with n=100 larvae per sample. (B) Genomic binding localization of average Dam:KDM5 TaDa profile (generated from four normalized replicates) showing enhanced binding near the TSS. (C) Distribution of Dam:KDM5 binding genomic regions showing enrichment for promoter-proximal regions. (D) Gene Ontology Biological Process (GO-BP) analyses of candidate KDM5 target genes identified from Dam:KDM5 TaDa. Representative terms shown, full list in Table S3. (E) Representative genome browser image showing binding of KDM5 in prothoracic gland TaDa experiment juxtaposed with published data sets from whole adult KDM5 ChIP-seq and ganglion mother cell TaDa. (F) Venn diagram showing strong overlap of KDM5-bound genes in prothoracic gland cells, whole adults, and ganglion mother cells. Prothoracic Gland TaDa: Whole Adult ChIP bound gene overlap p<0.00001; Prothoracic Gland TaDa: GMC TaDa bound gene overlap p<0.00001 (Fisher’s exact test).

In total, KDM5 peaks within the prothoracic gland mapped to 5815 genes using a cutoff of false discovery rate (FDR) < 0.01 (Table S1). Gene Ontology (GO) analyses for Biological Processes enriched in this gene list produced a range of terms including processes related to cellular transport, metabolism, and signaling (Fig. 3D, Table S3). To assess the KDM5 binding targets in the prothoracic gland in relation to other contexts, we compared these data to existing ChIP-seq and TaDa data sets from whole adult flies and ganglion mother cells (neuronal precursors), respectively (Hatch et al., 2021, Liu and Secombe, 2015) (Fig. 3E). This revealed a highly significant overlap via comparison between the prothoracic gland TaDa and either data set as well as a total of 2463 genes that were bound in all three data sets (42.3% of all prothoracic gland targets) (Fig. 3F). This overlap of KDM5 targets may represent genes regulated by KDM5 across developmental stages and tissues. Overall, KDM5 appears to have the potential to regulate a large portion of the coding genome in the prothoracic gland, and these data are consistent with KDM5 having both tissue-specific functions and functions that are common across cell types.

To determine the functional relationship between KDM5 binding and target gene expression in the prothoracic gland, we performed bulk RNA-seq on dissected ring glands of wild-type and *kdm5^140^* wandering 3^rd^ instar larvae. Similar to previous transcriptional studies, mRNA-seq was carried out from dissected ring glands to assay the prothoracic gland transcriptome, as this cell type comprises the majority of the mass of the ring gland (Uryu et al., 2018, Christesen et al., 2017, Ou et al., 2016, Di Cara and King-Jones, 2016, Nakaoka et al., 2017). Using a stringent cutoff of FDR < 0.01, we identified 2424 differentially expressed genes (DEGs) in *kdm5^140^* ring glands, 1276 of which were downregulated and 1148 that were upregulated (Fig. 4A, Table S2). To determine which genes were likely to be directly regulated by KDM5, we integrated this gene expression data with the genomic binding data. 1290 (53.2%) of the *kdm5^140^* DEGs had an associated KDM5 promoter peak based on the prothoracic gland TaDa data (Fig. 4A, B). As seen in previous *kdm5* mutant RNA-seq experiments, direct KDM5 targets exhibit relatively subtle changes to gene expression (downregulated direct DEG log2FC (log2 Fold Change) average = −0.660, upregulated = 0.910), and the DEGs with the largest log2FC appear to be indirectly regulated by KDM5 (Fig. 4A) (Hatch et al., 2021, Belalcazar et al., 2021, Liu and Secombe, 2015, Zamurrad et al., 2018). Gene Ontology (GO) analyses of the full list of DEGs produced primarily metabolic terms, including biological processes involving mitochondria and lipid metabolism (Fig. 4C, Table S3). The enrichment for these terms appeared to be driven by downregulated DEGs, as analysis of that subset produced many of the same GO terms, while upregulated genes featured GO biological processes involving cellular transport and chromatin dynamics (Fig. 4C’-C’’, Table S3). Among the direct DEGs, there was a similar trend with the top GO analysis terms related to mitochondrial processes and cellular respiration (Fig. 4D-D’’, Table S3). Taken together, these genome binding and transcriptomic analyses reveal that gene expression programs under the direct regulation of KDM5 span various cellular processes in the prothoracic gland, particularly those involving metabolism and mitochondria.

**Figure 4:**
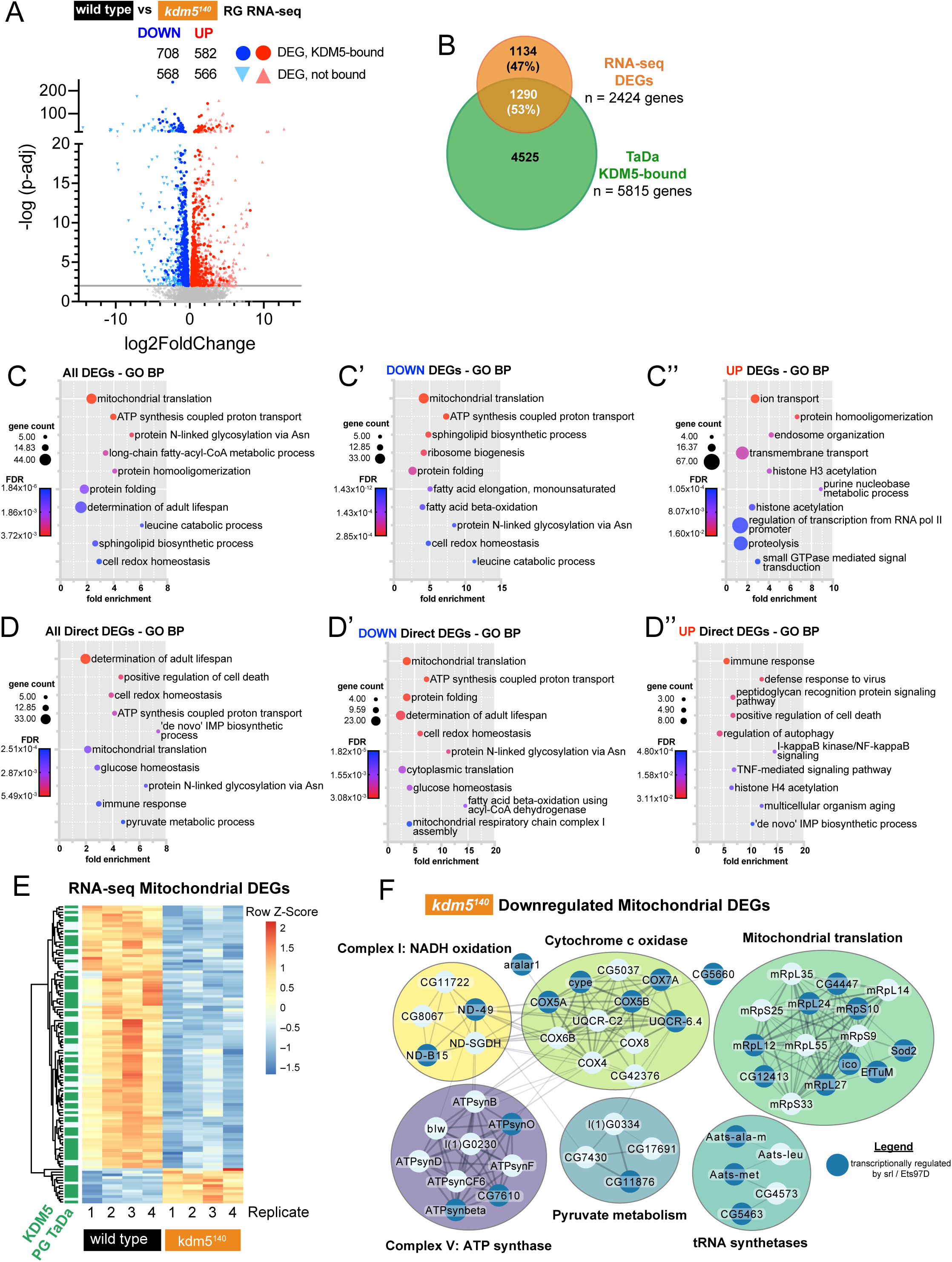
Changes to the transcriptome via bulk RNA-seq reveals transcriptional dysregulation of mitochondrial genes in *kdm5^140^*mutants. (A) Volcano plot of differentially expressed genes (DEGs) between *kdm5^140^*and wild-type wandering 3^rd^ instar larval ring glands. Genes with a false discovery rate (FDR) < 0.01 are colored blue (downregulated) and red (upregulated), and those directly bound in KDM5 TaDa are highlighted as bolded circles. RNA-seq was performed in quadruplicate for each genotype with n=80 ring glands per sample. (B) Venn diagram showing overlap of DEGs in *kdm5^140^* ring glands and direct KDM5 targets identified in prothoracic gland TaDa. (C-C’’) Gene Ontology Biological Process (GO-BP) analyses of DEGs in *kdm5^140^* ring glands. All DEGs (B), downregulated DEGs (B’), and upregulated DEGs (B’’) subsets were analyzed using GO DAVID. Representative terms shown, full lists in Table S3. (D-D’’) Gene Ontology Biological Process (GO-BP) analyses of DEGs that were directly bound in Dam:KDM5 prothoracic gland TaDa. All direct DEGs (B), downregulated direct DEGs (B’), and upregulated direct DEGs (B’’) subsets were analyzed using GO DAVID. Representative terms shown, full lists in Table S3. (E) Heatmap showing RNA-seq FPKM (Fragments Per Kilobase of transcript per Million mapped reads) of 111 genes from the mitochondrion GO term that were differentially expressed (FDR <0.01) in *kdm5^140^* ring glands. KDM5-bound genes in prothoracic gland TaDa are annotated in green in the column on the left side. (F) Physical protein interaction networks of mitochondrial genes downregulated in *kdm5^140^* ring glands. Genes potentially regulated by both KDM5 and srl/Ets97D (from microarray data in Tiefenbock et al. 2010)) are highlighted with darker blue nodes. Single nodes without physical connection edges excluded from image. Created with Cytoscape.

### KDM5-regulated transcription is developmentally required for proper mitochondrial dynamics

Our genome-wide analyses revealed that the transcriptional changes caused by the dysregulation of KDM5-mediated mechanisms in *kdm5^140^* mutants particularly affected mitochondria-related genes in the prothoracic gland. In addition to *Drosophila*, KDM5 proteins have been previously associated with mitochondrial activity in mammals and humans, although the mechanisms and biological implications of these connections remain unclear (Liu and Secombe, 2015, Varaljai et al., 2015, Liu et al., 2023). Within the Gene Ontology database, 353 *Drosophila* genes are classified in the mitochondrion biological processes category, and of these, 111 genes were differentially expressed in *kdm5^140^* animals (FDR < 0.01). Consistent with the GO analyses, the majority of these mitochondrial genes both showed downregulated expression across our RNA-seq replicates in *kdm5^140^* compared to wild type and were directly bound in the prothoracic gland TaDa data set (Fig. 4E). Investigation of known physical interactions within this downregulated mitochondrial gene set identified connections including components of Cytochrome c oxidase and ATP synthase complexes, as well as mitochondrial translation (Fig. 4F). This transcriptomic data suggests that a key role for KDM5 may be to maintain the expression of genes critical to mitochondrial biology, and this could contribute to its essential developmental activities.

The large size of the polyploid prothoracic gland cells demands significant metabolic requirements to fuel the cellular processes contained within, and thus these cells may be particularly sensitive to perturbations in mitochondrial activity. In addition to generating key cellular metabolites, mitochondria in the prothoracic gland are important sites for Halloween gene (ecdysone hormone biosynthetic enzymes) activity in processing stored lipid precursors for hormone production (Sandoval et al., 2014, Jacobs et al., 2020, Pan et al., 2020). To test whether the gene expression changes associated with mitochondrial function were linked to the lethality caused by loss of *kdm5*, we sought genetic approaches aimed at attenuating this deficit. spargel (srl) and Ets97D (Delg), homologous to mammalian PGC1-α and GABPα/NRF-2, respectively, are known transcriptional activators of genes required for mitochondrial biosynthesis in *Drosophila* (Tiefenbock et al., 2010, Tain et al., 2017, Sainz de la Maza et al., 2022). Previously published microarray experiments showed that both proteins can activate many of the mitochondrial genes found to be downregulated in *kdm5^140^* animals (Fig. 4F, highlighted in darker blue) (Tiefenbock et al., 2010). In light of this transcriptional data, we tested whether transgenic expression of srl or Ets97D in the prothoracic gland could restore viability to *kdm5^140^* animals. While *spok*-Gal4-driven expression of srl failed to suppress *kdm5^140^*lethality, significantly, expression of Ets97D did restore viability and produce adult flies (Fig. 5A, B’’). This result provides evidence that the activation of mitochondrial function genes that are necessary for animal survival may be mediated by KDM5.

**Figure 5:**
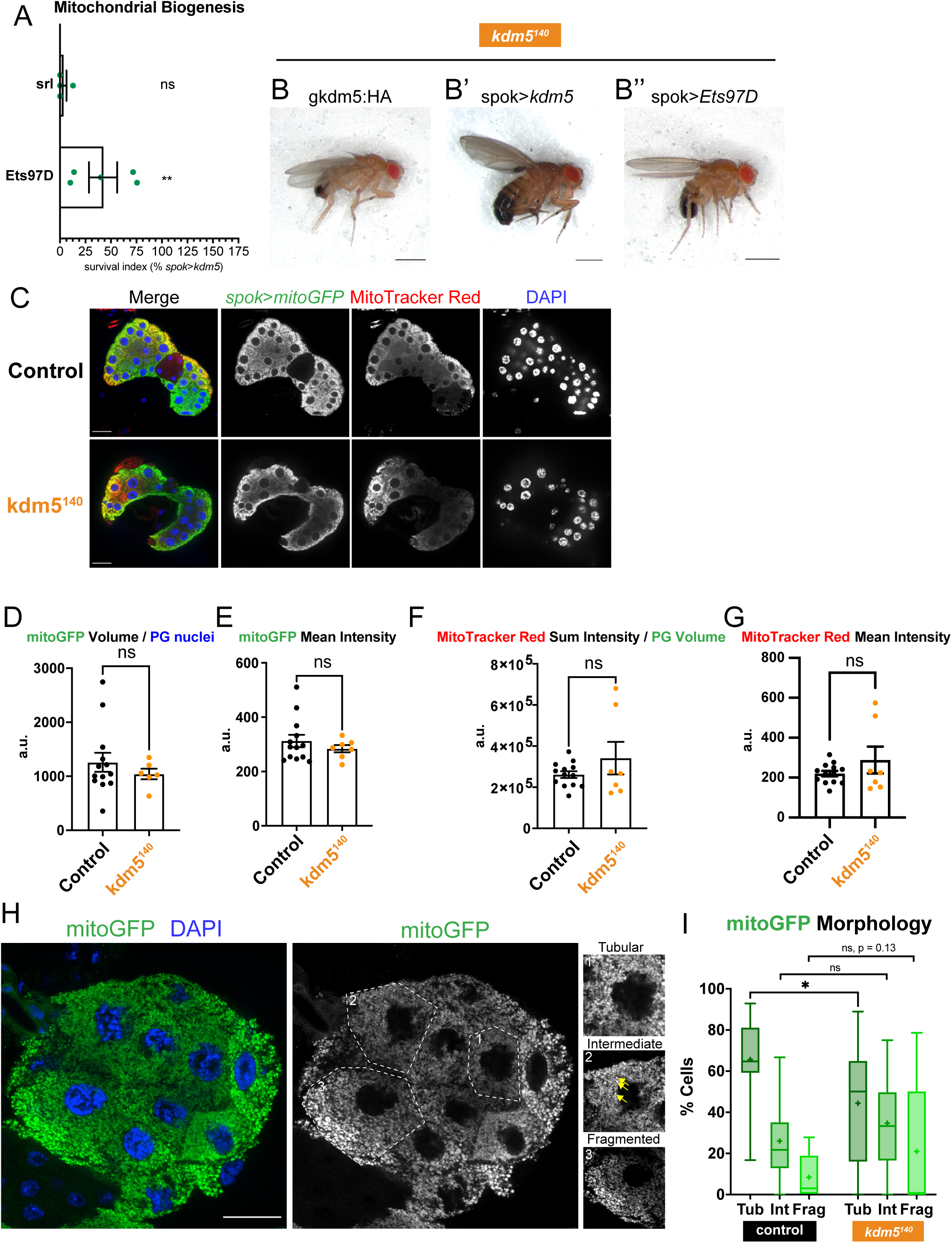
KDM5 regulates mitochondrial dynamics in the prothoracic gland that are critical for development. (A) Quantification of survival index for expression of mitochondrial biogenesis factors in *kdm5^140^* background relative to *spok*-Gal4>UAS-*kdm5*. n = 757-922 (mean n = 840) per genotype tested. **p<0.01; ns, not significant (Fisher’s exact test compared to no UAS control). Error bars: mean + s.e.m. (B-B’’) Representative images of *kdm5^140^* adult flies with lethality suppressed by genomic region *kdm5:HA* transgene (B), *spok*>*kdm5* (B’), and *spok*>*Ets97D* (B’’). Scale bars represent 750 μm. (C) Representative images (single Z slices) of larval ring glands expressing *spok*>*mitoGFP* and stained for GFP and MitoTracker Red. Scale bars represent 20 μm. Control genotype is *kdm5^140^*/CyO-GFP heterozygous internal control animals. (D) Quantification of total mitoGFP signal volume in each prothoracic gland normalized by number of nuclei in that sample. a.u. = arbitrary units. n = 7-13 per genotype tested. ns, not significant. (Wilcoxon rank sum test). Error bars: mean + s.e.m. (E) Quantification of mean mitoGFP signal intensity across each prothoracic gland. a.u. = arbitrary units. n = 7-13 per genotype tested. ns, not significant. (Wilcoxon rank sum test). Error bars: mean + s.e.m. (F) Quantification of total MitoTracker Red signal sum intensity in each prothoracic gland normalized by number of nuclei in that sample. a.u. = arbitrary units. n = 7-13 per genotype tested. ns, not significant. (Wilcoxon rank sum test). Error bars: mean + s.e.m. (G) Quantification of mean MitoTracker Red signal intensity across each prothoracic gland. a.u. = arbitrary units. n = 7-13 per genotype tested. ns, not significant. (Wilcoxon rank sum test). Error bars: mean + s.e.m. (H) Representative image (single Z slice) of larval ring gland expressing *spok*>mitoGFP and stained for GFP. Insets demonstrate representative cells of each morphological classification. Yellow arrows indicate fragmented mitochondria within example Intermediate morphological cell. Scale bars represent 20 μm. (I) Quantification of mitoGFP morphological classifications normalized to number of cells quantified per sample. n = 9-17. *p<0.05; ns, not significant (nonparametric unpaired t test). Error bars: mean + s.e.m.

To assess whether *kdm5^140^*animals exhibited visible mitochondrial phenotypes, we expressed a UAS-*mitoGFP* reporter with *spok*-Gal4 to examine mitochondrial networks in prothoracic gland cells (Fig. 5C). Assaying overall mitochondrial mass by quantifying the mitoGFP signal volume and mean intensity per cell revealed no differences between *kdm5^140^* and control animals, indicating no change in mitochondrial abundance (Fig. 5D, E). To assess mitochondrial energetics, we stained with MitoTracker Red, a reagent that is retained in the mitochondrial matrix of active mitochondria where the membrane is hyperpolarized (Wong et al., 2020). Similar to mitoGFP, the MitoTracker Red signal showed no significant changes at a tissue-wide level in terms of sum intensity per prothoracic gland cell nor mean intensity in *kdm5^140^*animals compared to controls (Fig. 5F, G). Focusing our analysis to the cellular scale, we examined the morphology of the mitoGFP-marked mitochondrial networks, defining them as tubular, fragmented, and intermediate, as has been done in previous studies (Fig. 5H) (Deng et al., 2015, Kashatus et al., 2015). Quantifying the cellular proportion of each morphological category, prothoracic glands from control animals display a majority of tubular cells with elongated and highly branched mitochondria (Fig. 5I). In contrast, *kdm5^140^* prothoracic glands showed a significant decrease in the proportion of tubular cells, with these glands featuring more rounded and isolated mitochondrial populations of the intermediate and fragmented categories. These results indicate that although there are no changes to overall abundance, mitochondrial biology is disrupted at the organelle level in *kdm5^140^* mutants. The increase in fragmented mitochondria in *kdm5^140^* could be due to defects in any of a number of mitochondrial dynamics including biosynthesis, fusion, or turnover or, alternatively, as a stress response to other cellular defects. Future analysis of specific mitochondrial components and bioenergetic processes as well as phenomena such as ROS (reactive oxygen species) and ER (endoplasmic reticulum) stresses will be fundamental in better understanding these *kdm5*-induced defects. Taken together, our data show that KDM5 transcriptional regulation in prothoracic gland cells is needed for mitochondrial homeostasis, and defects in mitochondria and cellular respiration in the prothoracic gland are key contributors to the lethality caused by loss of KDM5 (Fig. 6).

**Figure 6:**
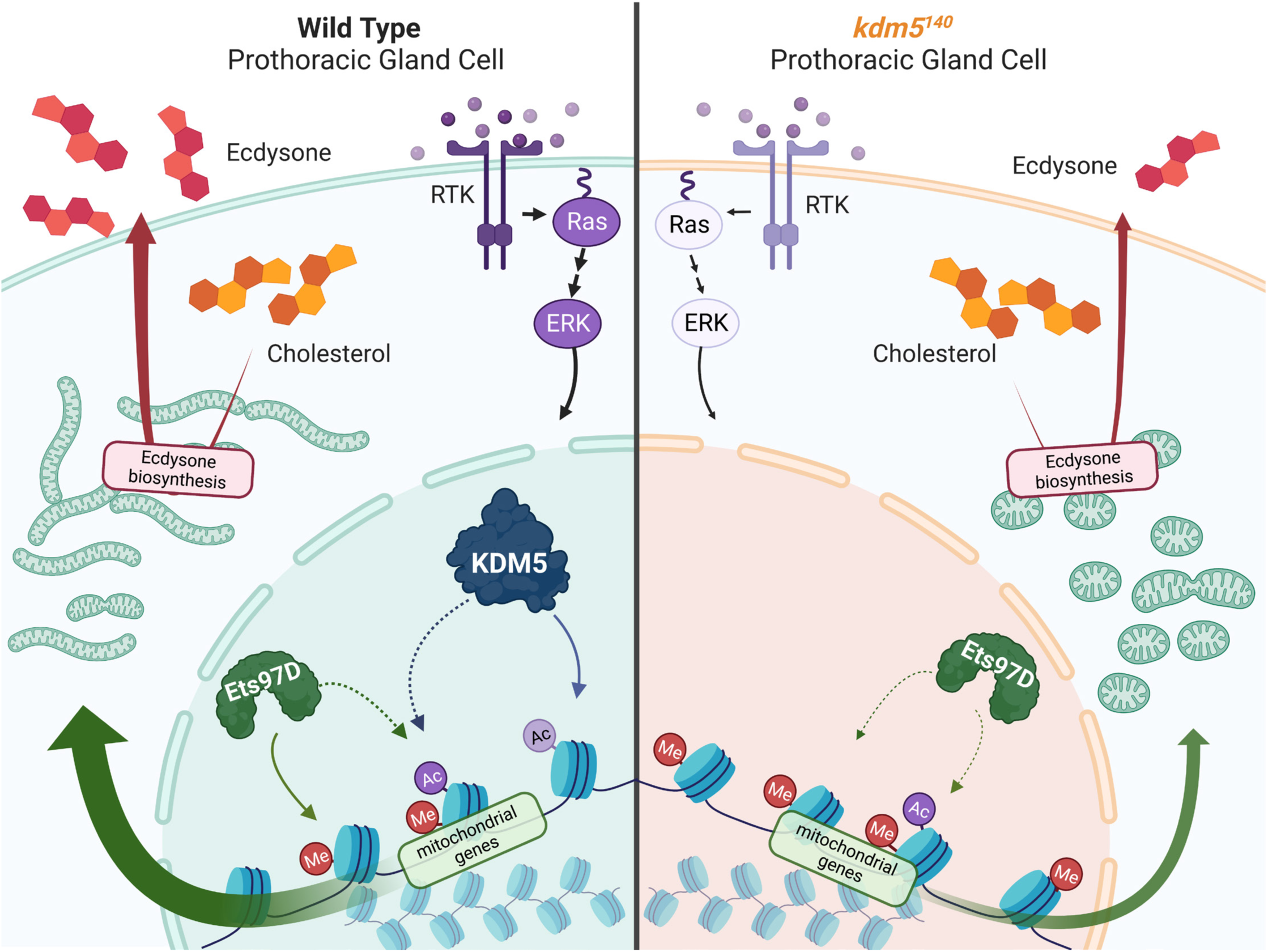
Model for KDM5-mediated transcriptional regulation of mitochondrial biology in prothoracic gland cells. (A) KDM5 regulates gene expression programs in the prothoracic gland coordinating proper MAPK signaling and mitochondrial morphology that are critical for development. Created with BioRender.

## DISCUSSION

In this study, we incorporated unbiased genome-wide data with targeted genetic and cellular analyses in order to expand our understanding of how KDM5 functions to regulate critical cellular processes during development. While expression and phenotype data show that KDM5 is important across many cell types, we focused this study on the prothoracic gland, where we have demonstrated that KDM5-regulated expression programs are important for survival (Drelon et al., 2019). This work has revealed important roles for KDM5 with respect to intracellular signaling and processes, notably MAPK and mitochondrial homeostasis. Consistent with our prior observation that loss of KDM5 resulted in reduced MAPK signaling, prothoracic gland-specific expression of MAPK-activating RTKs or ERK suppressed *kdm5^140^* (null) lethality (Drelon et al., 2019). Despite the energy-regulatory pathway of autophagy being one of the best characterized cellular processes downstream of signaling pathways in the prothoracic gland, enhancing or attenuating this process had no effect on the lethality of *kdm5^140^*animals. Instead, our KDM5 genomic binding and gene expression analyses point to a vital role for KDM5 in the regulation of a range of metabolic processes needed for cellular homeostasis, particularly mitochondrial function. Confirming the importance of KDM5-regulated expression programs that support mitochondrial activity, we find morphological changes to these organelles. Moreover, these changes are likely to be important for KDM5’s essential functions, as expression of the transcription factor Ets97D/GABPα, a known regulator of genes needed for mitochondrial function, suppressed *kdm5^140^* lethality.

This is not the first study to find an association between KDM5 proteins and the regulation of mitochondrial genes, which we have observed previously in adult flies, and others have seen with human KDM5A/RBP2 and KDM5C (Lopez-Bigas et al., 2008, Liu and Secombe, 2015, Liu et al., 2023, Varaljai et al., 2015, Kim et al., 2022). Muscle cells of adult flies harboring a hypomorphic combination of *kdm5* alleles showed abnormal mitochondrial shape, altered expression of redox-related genes, and increased sensitivity to oxidative stress (Liu et al., 2014, Liu and Secombe, 2015). Interestingly, most of the genes found to be altered in adult flies linked to altered cellular redox state do not overlap with those observed here in the prothoracic gland that were linked to respiratory chain complexes and translation. KDM5 is therefore likely to play different roles in distinct cell types. In human cells, the KDM5-mitochondria relationship has primarily been examined during the process of differentiation. While we observed KDM5 to be required for the activation of mitochondrial gene expression, in promonocytic (monocyte and macrophage precursors) and myogenic precursor cells, KDM5A represses mitochondrial genes (Lopez-Bigas et al., 2008, Varaljai et al., 2015). Consistent with the disparate changes to transcription, loss of *Drosophila* KDM5 and inhibition of human KDM5A led to distinct changes to mitochondrial morphology. Reducing KDM5A led to more dense tubular mitochondrial networks, while we observe mitochondrial fragmentation in prothoracic gland cells lacking all KDM5 function. In findings more similar with our data, KDM5C-deficient monocytes and osteoclasts have decreased mitochondrial gene expression resulting in decreased bioenergetic metabolism (Liu et al., 2023). Therefore, KDM5 proteins regulate the transcription of genes integral to mitochondrial function, but it is possible that whether this results in increased or decreased expression depends on the energy demands of a given cell type and/or the developmental cellular context. Indeed, it is notable that in muscle cell differentiation, KDM5A appears to function as part of an E2F/DP/pRb axis to regulate mitochondrial function in myogenic precursor cells, while we find that E2F1/DP does not suppress *kdm5^140^* lethality (Varaljai et al., 2015). Integrating these studies, it is apparent that mitochondrial and other metabolic genes are conserved targets of KDM5-mediated transcriptional regulation across species. Considering the breadth of KDM5 targets in the prothoracic gland TaDa data as well as others, KDM5 appears to occupy a large number of loci across the genome and may be utilized by the cell in either activating or repressive mechanisms depending on the context of cellular conditions (Hatch et al., 2021, Liu and Secombe, 2015). Notably, prothoracic gland cells are terminally differentiated and polyploid, requiring different homeostatic dynamics than the differentiating precursor cells of the mammalian studies. This may lead to KDM5 interacting with distinct gene regulatory complexes, or possibly employing histone demethylase-dependent and independent activities to alter transcription. Indeed, based on our observation that flies lacking KDM5-mediated histone demethylase activity are viable, the regulation of mitochondrial-related genes in the prothoracic gland is expected to be independent of its enzymatic function (Drelon et al., 2019).

The transcription factors srl (ortholog of mammalian PGC1α) and Ets97D (ortholog of mammalian GABPα) are involved in the activation of many of the same genes required for mitochondrial function that are regulated by KDM5 (Tiefenbock et al., 2010). While we observe robust suppression of *kdm5^140^*-mediated lethality by expression of Ets97D, srl expression failed to do the same. This may reflect differences in the function of these transcription activators, or based on studies of PGC1α, that post-translational modifications are particularly important for srl activity (Luo et al., 2019, Tain et al., 2017). While ectopic expression of Ets97D can compensate for the loss of KDM5 in the prothoracic gland, it is not clear whether *kdm5^140^* animals die due to reduced Ets97D activity, as expression of this gene was not altered in our RNA-seq data (Table S2). It remains possible that the level of Ets97D protein is altered by loss of KDM5, or that both Ets97D and KDM5 present at mitochondrial function genes promotes appropriate levels of gene activation. Defining the molecular details of the KDM5-Ets97D-mitochondrial pathway will require additional genetic and cell biological analyses.

The simplest synthesis of our *kdm5^140^* suppression experiments is that KDM5 is needed for proper activation of the MAPK pathway and that this alters the activation of genes related to mitochondrial function, possibly through Ets97D. Similar to Ets97D, our RNA-seq data did not reveal significant changes to components of the MAPK pathway, thus it remains unknown precisely how KDM5 leads to altered signaling. The MAPK cascade inputs into many processes across the cell, impacting metabolism through a variety of levels of regulation. While the relationships between MAPK and metabolic processes such as autophagy and glycolysis are more established in the literature, some studies have found direct connections between MAPK signals and mitochondrial biology (Kashatus et al., 2015, Javadov et al., 2014, Galli et al., 2009, Haq et al., 2013). In fact, most of the existing links between MAPK and mitochondria have been identified in the context of cancer cells and RASopathy developmental disorders. Mitochondrial dynamics can be altered in various cancers, and some studies have looked at mitochondria as a potential target to antagonize MAPK-driven tumors (Serasinghe et al., 2015, Marchetti et al., 2018, Ferraz et al., 2020, Corazao-Rozas et al., 2016). Furthermore, RASopathies, a collection of rare diseases driven by germline MAPK mutations, exhibit forms of mitochondrial dysfunction that contribute to bioenergetic defects (Kontaridis and Chennappan, 2022, Dard et al., 2018). In both instances of cancer and developmental disorders, KDM5 proteins may be involved in the regulation of this axis of MAPK-mediated metabolic changes. The potential role for KDM5 with both MAPK signaling and mitochondrial regulation indicates that there’s potential to consider KDM5 when treating these disorders.

One outstanding question in these studies of *kdm5*-induced lethality is what roles KDM5-mediated transcriptional programs play specifically within the prothoracic gland that are sufficient for these cells to rescue lethality at an organismal level. Anoar et al. (2021) hypothesize that neurons are particularly susceptible to mitochondrial defects because of high energetic demands and because as long-lived post-mitotic cells, they cannot dilute out defective organelles by cell division (Anoar et al., 2021). Similarly, prothoracic gland cells exit the cell cycle in the embryonic stage and must survive as large, polyploid cells with high bioenergetic requirements into pupal stages to coordinate the *Drosophila* developmental programs. KDM5-mediated mitochondrial regulation may be a key facet in the life cycle of the prothoracic gland cells in maintaining metabolic homeostasis as the cells undergo endoreplication and regulated production of the steroid hormone ecdysone. While these data suggest that raising *kdm5^140^* animals on food supplemented with ecdysone should suppress their lethality, this is not the case (Drelon et al., 2019). It is possible that ecdysone-supplemented *kdm5^140^* animals fail to consume enough ecdysone to progress completely through pupal stages during which they must subsist entirely off stored nutrients. However, *kdm5^140^* animals are able to undergo metamorphosis and form adult structures when raised on standard food and therefore must have sufficient prothoracic gland capabilities to generate the large final metamorphic pulse of hormone. Alternatively, while *kdm5^140^* animals are able to stimulate gross adult structure formation, some of the finer details of the underlying tissue, particularly synapse formation between neurons in the brain and into peripheral tissues, may depend on not just the quantity of ecdysone hormone but also the specific timing of ecdysone pulses. During metamorphosis, the neuronal networks across the animal undergo significant growth, pruning, and synapse formation for innervation across the newly formed adult body (Truman and Riddiford, 2023). This neuronal patterning is coordinated in part by ecdysone-responsive transcriptional elements, and likely hinges on proper timing for synaptic inputs and outputs to meet appropriately. Overcoming the *kdm5*-dependent defects by transgene-mediated modulation of mitochondrial dynamics may restore prothoracic gland cell homeostasis and function sufficiently for the ecdysone production and release program to successfully guide this neuronal remodeling that needs to occur in pupae. Future studies analyzing the relationship between KDM5-regulated mechanisms, ecdysone temporal dynamics, and mitochondrial homeostasis in the prothoracic gland will be key in defining these essential developmental programs.

## MATERIALS AND METHODS

### Fly husbandry

All flies were kept at 25°C on standard food at 50% humidity and a 12 hour light/dark cycle unless otherwise stated. Food (per liter) contained 18 g yeast, 22 g molasses, 80 g malt extract, 9 g agar, 65 g cornmeal, 2.3 g methyl para-benzoic acid and 6.35 mL propionic acid. For studies comparing wild-type and *kdm5^140^* mutant larvae, animals were matched for developmental stage, not chronological age, as we have done previously (Belalcazar et al., 2021, Drelon et al., 2018, Drelon et al., 2019, Hatch et al., 2021). Thus, at 25°C, control wandering 3^rd^ instar larvae were collected ~120 hours after egg laying (AEL), while *kdm5^140^* larvae were collected ~168 hours AEL. For all analyses, we used equal numbers of male and female animals and pooled data since we did not observe any sex-specific effects. In all experiments testing suppression of *kdm5^140^* lethality, vials resulting in n<12 or n>80 total eclosed adult flies were excluded from final analyses. This vial density was experimentally determined to be optimal for potential survival of *kdm5^140^*animals as under- or overcrowding outside this density introduced additional variables, including inconsistent food conditions and larval competition with control CyO-GFP (heterozygous) animals.

### Fly strains and genetics

A detailed list of the genotypes of the flies used in each figure is included in the Key Resources Table in the Appendix.

The *kdm5^140^* mutant allele, *kdm5:3xHA*, *UASp-kdm5:HA*, *UAS-LT3-dam:kdm5*, and genomic region *kdm5:HA* transgenes have previously been described (Drelon et al., 2018, Hatch et al., 2021, Navarro-Costa et al., 2016). The *spok*-Gal4, UAS-*torso*, UAS-*Alk*, and UAS-*Alk^CA^*lines were kindly shared by Michael O’Connor (U. Minnesota). The UAS-*srl* line was kindly shared by Grace Zhai (U. Miami) with permission from Christian Frei (U. Zurich). The UAS-*Ets97D* line was kindly shared by Martine Simonelig (Institut de Genetique Humaine) with permission from Christian Frei. The UAS-*LT3-dam* line was kindly shared by Andrea Brand (U. Cambridge, Gurdon). All other strains were obtained from the Bloomington *Drosophila* Stock Center (see Key Resources Table in the Appendix).

### Immunohistochemistry

Wandering 3^rd^ instar larval brain-ring gland complexes were dissected in ice cold 1X phosphate buffered saline (PBS) and fixed in 4% paraformaldehyde (PFA) in PBS at room temperature for 20 min. Samples were washed three times in 1X PBST (PBS + 0.1% Triton) for 10 min each. Brain-ring gland complexes were transferred to 0.5 μL tubes for blocking in 1X PBST + 5% normal donkey serum (NDS) for 30 min, followed by primary antibody incubation overnight while rotating at 4°C. After three 15 min washes in 1X PBST, samples were incubated in secondary antibodies at room temperature rotating for 2 hours. Samples were then washed three times in 1X PBST and ring glands were dissected from brain tissue in ice cold 1X PBS. Finally, ring glands were mounted with Fluoromount-G DAPI (Southern Biotech), and slides were stored at 4°C for imaging within 1-3 days.

A similar protocol was followed for mitochondrial immunostaining with the following exceptions. Larval brain-ring gland complexes were dissected in ice cold 1X Schneider’s Medium (Gibco, Thermo Fisher Scientific), and then incubated in 500 nM MitoTracker Red CMXRos (Invitrogen, diluted in 1X Schneider’s Medium) for 30 min protected from light. After two 1X PBS washes, samples were fixed in 4% PFA in PBS. Additionally, after secondary antibody incubation, samples were washed five times in 1X PBST prior to mounting.

The following primary antibodies were used: mouse anti-HA (1:100, Cell Signaling Technology) and rabbit anti-GFP (1:100, Invitrogen). Primary antibodies were prepared in 5% NDS/PBST. The following secondary antibodies were used: goat anti-mouse Alexa-568 (1:500, Thermo Fisher Scientific) and goat anti-rabbit Alexa-488 (1:500, Thermo Fisher Scientific). Secondary antibodies were prepared in 5% NDS/PBST.

### Image Acquisition and Processing

Images of prothoracic gland signaling pathway, pupal brain, and the model of KDM5 function in the prothoracic gland were created with BioRender.com. All tissue images were taken on a Nikon CSU-W1 Spinning Disk confocal microscope using a 100X immersion lens (NA = 1.45 oil) and 0.2 um Z-step size. Adult fly images were obtained using a stereomicroscope Carl Zeiss Stereo Discovery V12 with 12.5X magnification and captured using AxioVision Release 4.8 software. All images were processed with ImageJ. All Venn diagrams were generated using the R package BioVenn (v1.1.3) (Hulsen et al., 2008). Figures were composed using Adobe Illustrator.

### *kdm5^140^* Lethality Suppression Experiments

To identify signaling pathway components that suppressed *kdm5^140^* lethality, *kdm5^140^* /CyO-GFP; *spok*-Gal4 flies were crossed with *kdm5^140^* flies carrying a UAS transgene and allowed to lay eggs for 48 hours at 25°C. Animals were kept at 25°C, and all eclosed adults were scored. Using Mendelian ratios, we estimated the number of *kdm5^140^* animals expected in each cross based on the total internal control (CyO-GFP) adults eclosed as done previously (Drelon et al., 2019). The survival index was calculated as a percentage of the total viable (lethality-suppressed) *kdm5^140^* adults eclosed over the estimated number of *kdm5^140^* animals in the cross. Graphed survival index data points represent biological replicate crosses normalized to the positive control *spok*>*kdm5* rescue.

### Western Blotting

For each sample, three male and three female adult heads (age 1-3 days) were homogenized in PBS, denatured in 1X loading buffer (3X Laemmli sample buffer containing 187.5 mM Tris, 6% SDS, 30% glycerol, 0.03% bromophenol blue, and 10% β-mercaptoethanol) at 95°C for 5 min, run on a 6% 1.5 mm gel, and transferred to a PVDF membrane. The following primary antibodies were used: mouse anti-HA (1:2000, Cell Signaling Technology) and mouse anti-αTubulin (1:10000, DSHB). Secondary antibody used was rabbit anti-mouse (1:1000, Invitrogen). Blots were scanned and processed using Kwik Quant Imager (Kindle Biosciences) scanner.

### KDM5 Temporal Experiments

To identify the developmental windows requiring *kdm5* expression, *kdm5^140^*, *Ubi*-Gal4 / CyO-GFP flies were crossed with tub-Gal80^ts^, *kdm5^140^* / CyO-GFP; UAS*-kdm5:HA* flies and allowed to lay eggs for ~12 hours at either 18°C or 29°C. Animals raised at 18°C were transferred to 29°C to induce the expression of the *kdm5* transgene, and all eclosed adult flies were scored. Conversely, animals raised at 29°C were transferred to 18°C to repress the expression of the *kdm5* transgene, and adults were scored in the same way. For 18°C to 29°C shifts, days 1-15 were tested with, n > 100 flies eclosed for each day of shift. For 29°C to 18°C shifts, days 1-12 were tested in the same manner. The survival indices for these crosses were calculated in the same method as the *kdm5^140^* lethality suppression experiments. Graphed survival index data points represent vial replicates normalized to the positive control *Ubi*>*kdm5* at constant 29°C rescue.

### Targeted DamID and analyses

To profile the genomic regions bound by KDM5 in prothoracic gland cells, tub-Gal80^ts^; *spok*-Gal4 flies were crossed with flies carrying *UAS-LT3-dam* or *UAS-LT3-dam:kdm5* and allowed to lay eggs for 24 hours at 18°C. Animals were kept at 18°C for 5 days then transferred to 29°C for 2 days to induce the expression of the transgenes. Wandering 3^rd^ instar larvae were collected, flash frozen on dry ice, and stored at −80°C.

Tissue processing was performed as previously described with the following modifications (Marshall et al., 2016a). TaDa was performed in quadruplicate with replicates of 100 larvae that were homogenized and digested in Proteinase K in samples of 50 larvae then pooled into replicates of 100 larvae prior to DNA extraction. Larvae were homogenized in 75 uL UltraPure Distilled Water and 20 uL 500 mM EDTA then digested with Proteinase K for 1.5 hours. DNA extraction was performed using the Zymo Quick-DNA Miniprep Plus Kit. DpnI digestion, PCR adaptor ligation, DpnII digestion, and PCR amplification were performed as described. DNA was sonicated using a Diagenode Bioruptor Pico for 6 cycles (30 sec on/90 sec off at 4°C), and DNA fragments were analyzed using an Agilent Bioanalyzer to confirm ~300 bp fragment size. DamID adaptor removal and DNA cleanup were performed as previously described, and samples were submitted to BGI Genomics for library construction and sequencing.

Libraries were prepared at BGI Genomics following a ChIP-seq workflow. DNA fragments were first end-repaired and dA-tailed using End Repair and A-Tailing (ERAT) enzyme. Adaptors were then ligated for sequencing and ligated DNA purified using AMPure beads. DNA was then PCR amplified with BGI primers for 8 cycles and PCR purified with AMPure beads. DNA was then homogenized, circularized, digested, and again purified. DNA was then prepared into proprietary DNA nanoballs (DNB™) for sequencing on a DNBSEQ-G400 platform with 50 bp single-end read length and 20M clean reads passing filter.

For targeted DamID analyses, sequencing data were aligned to the Dm6 *Drosophila melanogaster* genome and processed using damidseq_pipeline as previously described (Marshall and Brand, 2015, Marshall et al., 2016a, Marshall and Brand, 2017). After converting to bedgraphs via damidseq_pipeline, peaks were called using find_peaks (using the parameters fdr = 0.01, min_quant = 0.9) on the averaged replicates, and genes overlapping peaks identified using peaks2genes (Marshall et al., 2016a, Marshall et al., 2016b).

For genome localization analyses, the R package ChIPseeker (v1.34.1) was used with the average KDM5 binding BED file to generate profiles (Wang et al., 2022). Gene Ontology (GO) enrichment analysis of KDM5 bound genes (FDR < 0.01) utilized GO DAVID database (v2021), specifically annotation GOTERM_BP_DIRECT (Sherman et al., 2022). Genome browser image was generated using pyGenomeTracks (v3.8) utilizing BedGraph or bigWig files from: adult fly KDM5 ChIP-seq (<u>SRX1084165</u>) and larval neuronal precursor KDM5 TaDa (<u>GSE166116</u>) (Lopez-Delisle et al., 2020).

### RNA sequencing

RNA sequencing (RNA-seq) was carried out on pooled ring glands dissected from control (*w^1118^*) and *kdm5^140^* wandering 3^rd^ instar larvae. Ring glands were dissected and washed three times in ice cold 1X PBS, transferred to TRIzol, flash frozen on dry ice, and stored at −80°C. 80 dissected ring glands were pooled to form each of the four replicates. Total RNA was isolated with TRIzol and Phasemaker tubes (Invitrogen), and quality was assessed by Agilent Bioanalyzer before sending to Novogene for library construction and sequencing. mRNA was purified from total RNA using poly-T oligo-attached magnetic beads. After mRNA fragmentation, first strand cDNA and second strand cDNA were synthesized, and cDNA fragments were purified with AMPure XP system to select for suitable sizes for PCR amplification. Library quality was assessed on the Agilent Bioanalyzer 2100 system. Libraries were sequenced on the Ilumina NovaSeq PE150 platform (2 x 150bp cycles). Alignment of raw reads to the reference genome (dm6) was performed using Hisat2 (v2.0.5) for mapping, assembly via StringTie (v1.3.3b), quantification via featureCounts (v1.5.0-p3), normalized, and differential expression was determined with the DESeq2 package (1.20.0) (Pertea et al., 2016, Love et al., 2014, Liao et al., 2013).

Gene Ontology enrichment analysis of protein-coding genes found to be dysregulated in *kdm5^140^*RNA-seq data (1% FDR cutoff) was carried out using GO DAVID annotation GOTERM_BP_DIRECT (Sherman et al., 2022). The heatmap was generated using the R package pheatmap (v1.0.12) (Kolde, 2012). Physical interaction networks were determined using String and visualized using Cytoscape (v3.9.1) (Shannon et al., 2003).

### Quantification and statistical analyses

All experiments were carried out in biological triplicate (minimum) and numbers (*n*) are provided for each experiment in the Figure Legends.

For *kdm5^140^* lethality suppression experiments, a Fisher’s exact test was performed in R Studio (v2023.03.0) comparing survival index of each genotype to the no UAS control genotype as done previously with ****p<0.0001, ***p<0.001, **p<0.01, *p<0.05; ns, not significant (Drelon et al., 2019, RStudio, 2020). For KDM5 binding Venn Diagram overlap, a Fisher’s exact test was performed in R Studio.

For mitoGFP and MitoTracker Red fluorescent images, the control genotype used was *kdm5^140^*/CyO-GFP heterozygous animals that developed from the same cross alongside the *kdm5^140^* homozygous animals because we have not seen the same developmental and lethality phenotypes from these animals (Drelon et al., 2019). Volocity software was used to quantify the intensity and 3-dimensional volume of the fluorescent signal in each channel. Student’s *t*-test comparing control and *kdm5^140^* genotypes was performed in GraphPad Prism (v9.5.1) (GraphPad, 2023). mitoGFP morphological quantifications were performed as follows. All images were blinded to genotype and analyzed at two Z-slice locations positioned 33% and 66% through the full Z-plane of the sample. At each Z-slice, all cells with nuclei clearly visible by DAPI signal at that Z-position were identified and classified for mitochondrial morphology of tubular, intermediate, or fragmented by scrolling through the Z-slices occupied by each identified cell. Tubular morphology consisted of zero visible fragmented round mitochondria, intermediate morphology consisted of primarily tubular morphology with >1 visible fragmented mitochondria, and fragmented morphology consisted primarily of fragmented mitochondria. The proportion of cells with each morphological classification was calculated per sample (individual prothoracic gland), and a parametric unpaired *t*-test was performed in GraphPad Prism comparing each morphological category between control and *kdm5^140^*animals.

## AUTHOR CONTRIBUTIONS

Conceptualization, M.F.R., J.S.; Methodology, M.F.R., O.J.M.; Investigation, M.F.R., O.J.M. and J.S.; Writing – original draft, M.F.R. and J.S., Writing – Reviewing and Editing, M.F.R., J.S., and O.J.M.; Funding acquisition, J.S. and O.J.M, Supervision, J.S. and O.J.M

## DECLARATIONS

### Ethics approval and consent to participate

N/A

### Availability of data and materials

KDM5 binding (TaDa) and gene expression (RNA-seq) data have been deposited in the Gene Expression Omnibus (GEO) under SuperSeries accession numbers GSE229077. Transgenic fly strains used in this research are available upon request to Julie Secombe.

## Acknowledgements

We thank members of the Secombe and Baker labs for their intellectual contributions to this project and comments on the manuscript. We also thank Andrea Briceno, Hillary Guzik, and Vera Desmarais of the Einstein Analytical Imaging Facility (AIF) for the confocal microscope training and technical assistance with capturing and quantifying images (NCI P30CA013330). We thank Melissa Fazari, Mimi Kim, Jaeun Choi, Kenny Ye, and Abdissa Negassa of the Einstein Statistics Consulting Center for assistance with experimental design and statistical analyses. We also appreciate the availability of stocks from the Bloomington *Drosophila* Stock Center (NIH P40OD018537) and are grateful to the Einstein Cancer Center Support Grant P30CA013330.

---

**Supplemental Figure 1:**
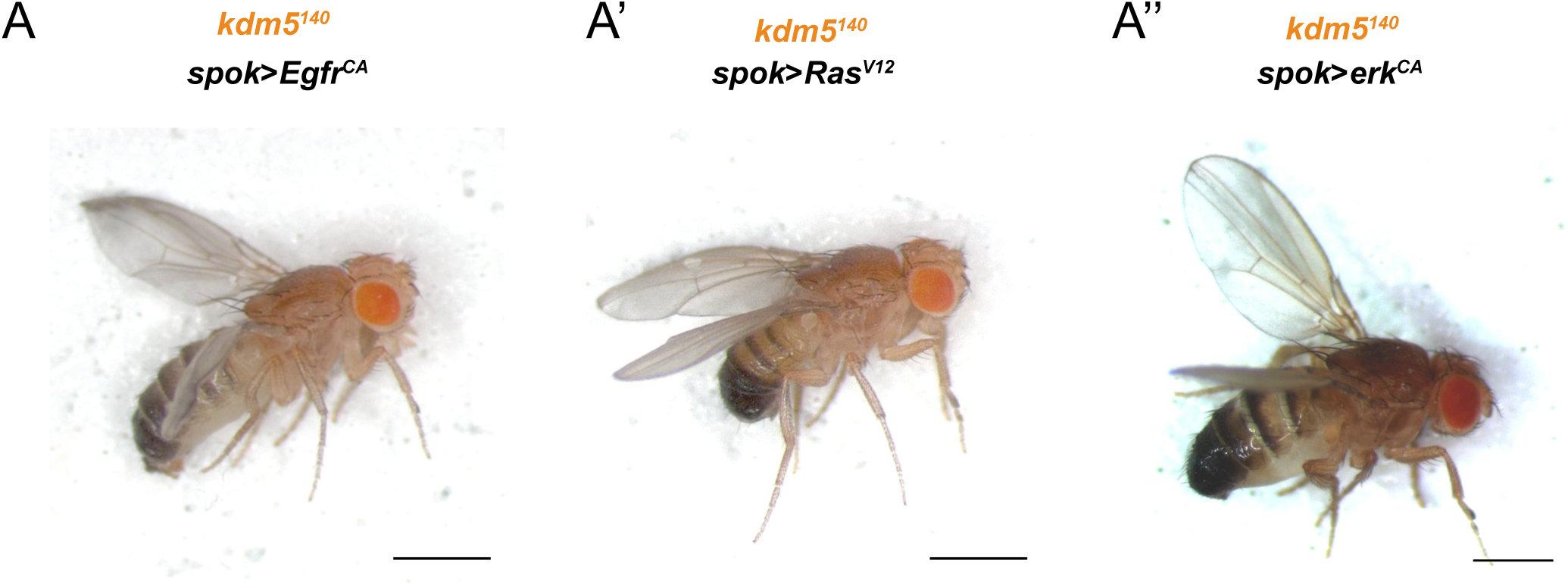
*kdm5^140^* adults with lethality suppressed by MAPK components. (A-A’’) Representative images of *kdm5^140^* adult fly with lethality suppressed by *spok*>*Egfr^CA^* (A, *spok*>*Ras^V12^* (A’), *spok*>*erk^CA^* (A’’). Scale bars represents 750 μm.

**Supplemental Figure 2:**
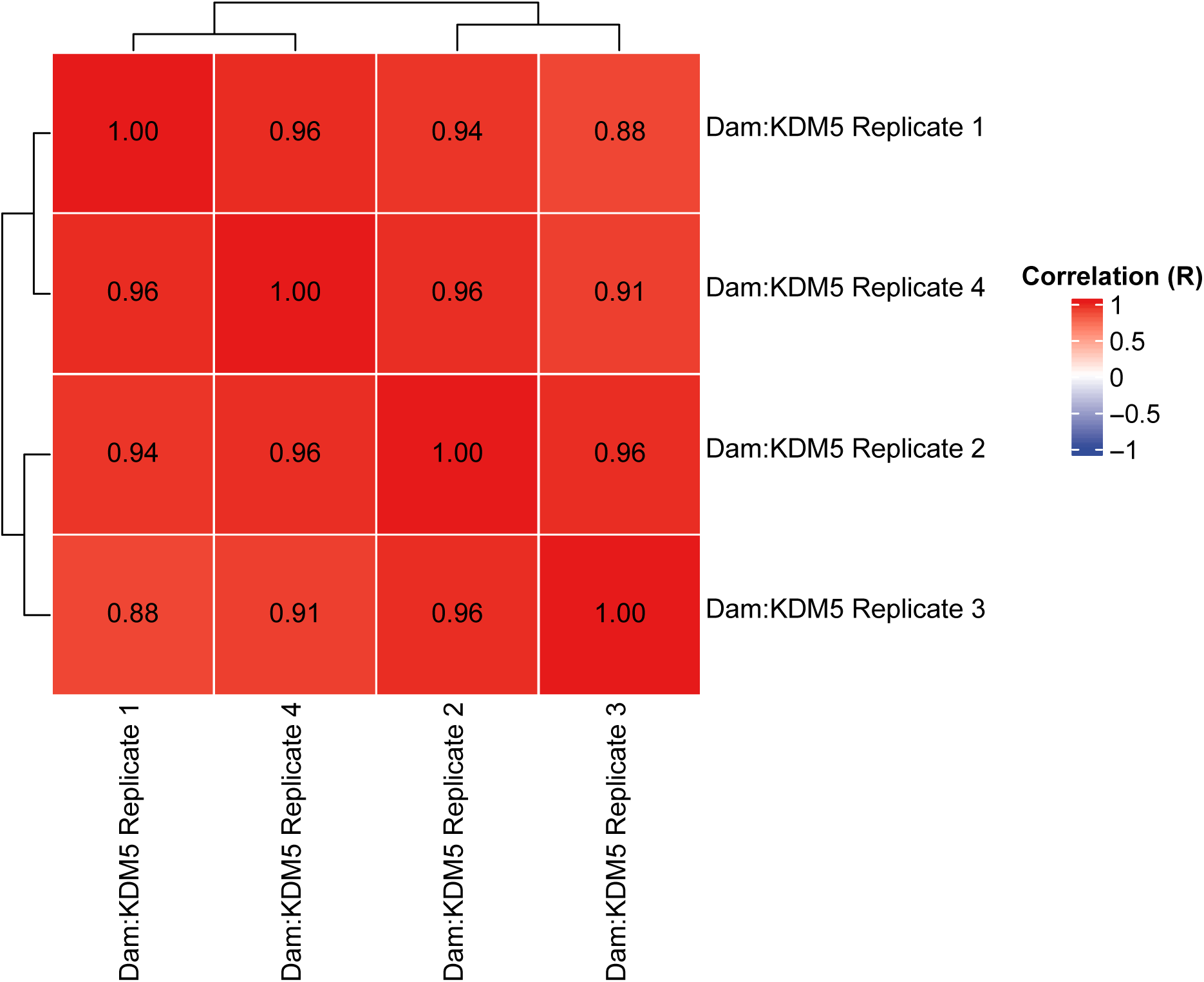
Targeted DamID replicate correlations. Plot showing correlation across binding profiles of Dam:KDM5 TaDa replicates.

**Supplemental Table S1: Targeted DamID-identified KDM5 target genes.**

List of Dam:KDM5-bound genes identified in Targeted DamID.

**Supplemental Table S2: RNA-seq analysis of *kdm5^140^* ring glands.**

wild type vs *kdm5^140^*ring gland RNA-seq data.

**Supplemental Table S3: Gene Ontology analyses of gene sets.**

Full lists of Gene Ontology (GO) terms generated via GO DAVID analyses.

